# Aging in a relativistic biological space-time

**DOI:** 10.1101/229161

**Authors:** D. Maestrini, D. Abler, V. Adhikarla, S. Armenian, S. Branciamore, N. Carlesso, Y-H. Kuo, G. Marcucci, P. Sahoo, R. Rockne

**Affiliations:** City of Hope, National Medical Center, Division of Mathematical Oncology; City of Hope, National Medical Center, Department of Pediatrics; City of Hope, National Medical Center, Department of Population Sciences; City of Hope, National Medical Center, Department of Diabetes Complications & Metabolism; City of Hope, National Medical Center, Department of Hematologic Malignancies Translational Science; City of Hope, National Medical Center, Gehr Family Center for Leukemia Research, City of Hope

## Abstract

Here we present a theoretical and mathematical perspective on the process of aging. We extend the concepts of physical space and time to an abstract, mathematically-defined space, which we associate with a concept of “biological space-time” in which biological clocks operate. We hypothesize that biological dynamics, represented as trajectories in biological space-time, may be used to model and study different rates of biological aging. As a consequence of this hypothesis, we show how the dilation or contraction of time resulting from accelerated or decelerated biological dynamics may be used to study precipitous or protracted aging. We show specific examples of how these principles may be used to model different rates of aging, with an emphasis on cancer in aging. We discuss the implications of this theory, including novel concepts that may improve our interpretation of biological aging.

## 1 Introduction

The connection between one’s chronological age and biological age is something that we all perceive. In a sense, it is the difference between the age you “feel” and the age you *are*. Some people look “young” for their age, while some become frail earlier than others [Ness et al., 2013]. Molecular “clocks” and markers of “biological age” can change throughout one’s lifetime. In some cases, the rate of change of a person’s biological age is *greater* than the rate of change of their chronological age. For instance, some cancers have been shown to increase biological aging, which can be described as an increase in the speed of biological clocks [Horvath, 2013].

From a mathematical perspective, biological clocks are frequently modeled as periodic or oscillating functions describing, for example, circadian rhythms [Klerman and Hilaire, 2007]. When studied in isolation, biological clocks can be described and predicted with periodic functions, which may speed up, slow down, or even stop and start over the course of a persons’ lifetime. The more complex case of multiple integrated biological clocks can be modeled with coupled oscillators and dynamical systems theory [Shiju and Sriram, 2017]. However, a fundamental assumption of these approaches is that the rate of change of these biological clocks are measured with respect to a *linear passage of time* and that the rate is independent from the biological *space* in which the biological clocks operate.

Here we investigate a mathematical model of biological space-time which includes the effects of time dilation and contraction resulting from accelerated or decelerating biological clocks, which may provide a new theoretical foundation and perspective on rates of aging. The principle assumption of our model is that a dynamic biological process may be represented as the motion of a (massless) point along a trajectory on a manifold. We then investigate the consequence of time dilation and contraction in terms of accelerating or decelerating motion of the point along the trajectory, and relate these concepts to rates of biological aging, with a particular focus on aging in cancer. In particular, we put forth the hypothesis that a biological space-time may be used to model aging from an arbitrary biological viewpoint relative to a common frame of reference. A consequence of this hypothesis is that precipitous (faster than chronological time) or protracted (slower) aging may be modeled by a dilation or contraction of time in the manifold in which the aging process occurs.

To the best of our knowledge, only a few groups have proposed similar concepts. Bailly and colleagues [Bailly et al., 2011, Longo and Montévil, 2014] have proposed a mathematical definition of “biological time” as a means to model biological rhythms and periodic biological processes. However, they do not define a biological “space” nor include the possibility of the dilation or contraction of time. Systems biology pioneer Denis Noble has proposed a theory of biological relativity, which asserts that there is no “privileged level of causation” in biology [Noble, 2012]. Nobel’s theory contends that biological processes occur on many scales in space and in time, and that these scales are coupled to each other and should not be separated; that no single scale is responsible for the dynamics of the whole. Noble’s theory implicitly couples space and time, but does not include a definition of biological space or concept of dilation or contraction of time. Consequently, our work is among the first to introduce both the mathematical interpretation and specific concept of a relativistic biological space-time to be used to study rates of biological aging.

This manuscript is structured as follows: first we describe the mathematical objects which we use to define biological space-time. Then we show how the dilation and contraction of time can be used to model precipitous or protracted aging. We then show examples of how these principles may be used to model aspects of the aging process with a particular emphasis on aging in cancer. We discuss the implications of this theory, and suggest novel biological quantities that may improve our interpretation of biological aging.

## 2 Biological space-time

Biology, and biological processes, are measured and observed in our conventional notion and understanding of physical space. Cells, tissues, and organisms move and change in a physical space that we can measure with length and time scales in conventional units. However, we may also consider the functional, or phenotype space in which biological processes can be represented as locations in the space. We refer to movement in a biological space as a sequence of locations in the space that form a trajectory. These general concepts have been used to characterize biological states such as hematopoietic differentiation, where 2- or 3-dimensional representations of biological space are constructed with dimension reduction techniques applied to high-dimensional single cell RNA-sequencing data [Nestorowa et al., 2016, Rizvi et al., 2017, Mojtahedi et al., 2016]. The idea of the *relativity* of aging is to apply the special relativity machinery to provide a rigorously defined mathematical framework to represent biological dynamics as trajectories on manifolds moving at different rates relative to a common frame of reference. In other words, the difference in aging between two different people will be explained in terms of different dynamics in biological space-time.

### 2.1 A manifold 𝔐 and submanifolds 𝔐*_i_*

In order to provide a conceptual picture of our mathematical framework, we imagine the life of a living person represented by a curve, or trajectory, on a connected smooth manifold. A manifold is a topological space that is locally, but not necessarily globally, Euclidean. The canonical example of a manifold is the Earth: locally flat, globally round. The complexity of all biological processes taking place in the body of the living person is then decomposed to single biological processes whose dynamics occur on a submanifold. In what follows we identify with the index *i* a possible biological process.

Given a subset *U_i_* of a topological space 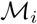 a *d_i_*-dimensional chart is an injective function *φ_i_* : *U_i_* ⊂ 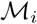 *→* ℝ*^di^*. A point *q_i_* on 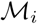 is identified by a set of *d_i_* spatial coordinates 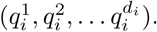 An atlas on 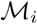 is the set 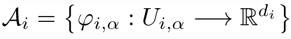 for some finite values of the index *α*, where the union *U*_*i*,1_ ∪ *U*_*i*,2_ ∪ … is the whole space 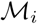. A space 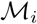 equipped with an atlas 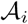 is a *d_i_*-dimensional differential manifold.

A trajectory *γ_i_* on 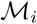 is a smooth map

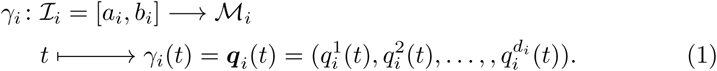

Here *t* ∈ [*a_i_,b_i_*] ⊂ ℝ is a parameter that is interpreted as the time variable associated to the manifold 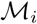. The curve *γ_i_* represents the time evolution of the *i*-th biological process starting from its beginning, *γ_i_(a_i_)*, to its end, *γ_i_(b_i_)* (Figure (1)).

**Figure 1:**
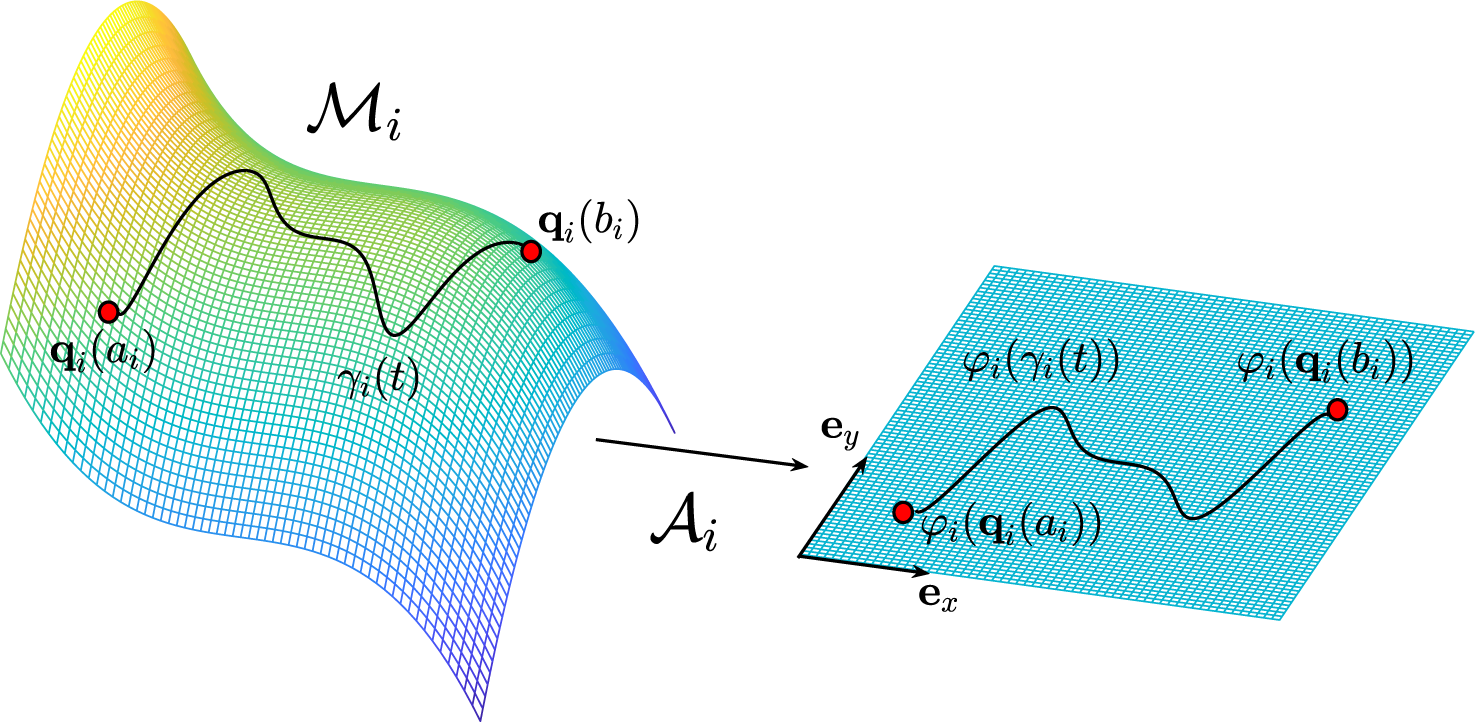
The trajectory *γ_i_* on the manifold 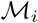; represents the evolution of the *i*-th biological process which starts at ***q**_i_(a_i_)* and ends at ***q**_i_(b_i_)*. The atlas 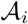 provides the connection with the Euclidean space ℝ^2^ identified by the two unit vectors *e_x_* and *e_y_*.

Given an interval 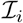 ⊂ ℝ, we now define the *i*-th biological *space-time* manifold 𝔐*_i_* as the *d_i_* + 1-dimensional Lorentzian manifold given by the Cartesian product

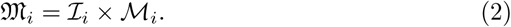

We assume there can be a countably infinite number of these processes and hence submanifolds. A point *𝒬_i_* on the manifold 𝔐*_i_* is then identified by a set of *d_i_* + 1 coordinates 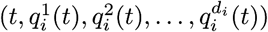. The invariant square of an infinitesimal line element between the points 𝒬*_i_* and 𝒬*_i_* + *d*𝒬*_i_*, referred to as *interval*, is then evaluated by

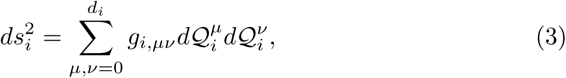

where *g_i,μν_* is the metric tensor of the *i*-th biological manifold whose entries are functions of the local coordinates, 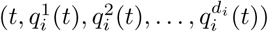. Knowledge of the metric tensor provides informations on the geometry of the manifold and vice-versa. It is therefore a key quantity which characterizes the biological space-time and it can be evaluated once the structure of the manifold is known or hypothesized. What we emphasize is that the notion of distance is not necessarily the usual Euclidean distance and therefore, when dealing with biological processes, distances should be measured only once the entries of the metric tensor are known.

We now consider the complexity of all biological processes by constructing the manifold

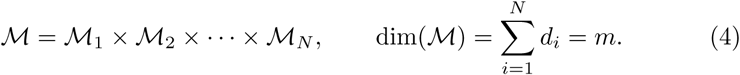

A point ***q*** on 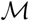 is then identified by the set of coordinates (***q*_1_, *q*_2_,…, *q_N_***) where each ***q***_i_ ∈ 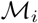, *i* = 1, 2,…, *N* and it represents a possible configuration, or state, in which the human body can be found. Following the definition given by Eq. (1), a trajectory Γ on 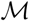 is a smooth map

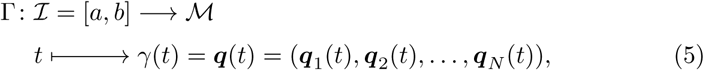

where *t* ∈ [*a,b*] ⊂ ℝ is a parameter that is interpreted as the time variable associated to the manifold 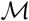 and the curve Γ represents the entire life of a person from its birth, Γ(*a*), to its death, Γ(*b*). The biological space-time 𝔐 for the whole body is then defined by the *m* + 1-dimensional Lorentzian manifold given by

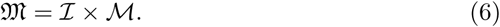

We consider the Cartesian product 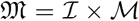 instead of 𝔐 = 𝔐_1_ × 𝔐_2_ × … × 𝔐*_N_* because in the latter case it is unclear how all time variables, one per each submanifold 𝔐*_i_*, will combine together and define the time on the resulting manifold 𝔐. A point 𝒬 on the manifold 𝔐 is then identified by a set of coordinates as follows

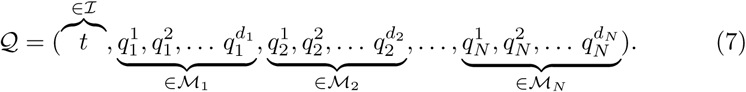

The idea for which time flows at different rates for different biological processes is now mathematically modeled by introducing a set of projection maps Π*_i_* such that

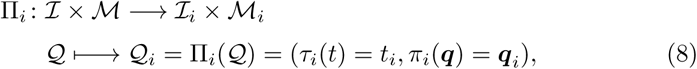

where the spatial part 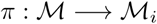 is the canonical projection map which maps the curve Γ on 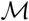 onto the curve *γ_i_*, on 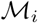 while the temporal part 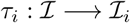 provides the connection between the time *t* on the manifold 𝔐 and the time *t_i_* on the *i*-th manifold 𝔐*_i_*, (Figure (2)). We emphasize that the relationship between *t* and *t_i_*, is not necessary linear. In fact, in the next section we will show that the time measured in a particular manifold depends upon the acceleration of the particle along a trajectory and it will produce a non linear relations between *t* and *t_i_*.

**Figure 2:**
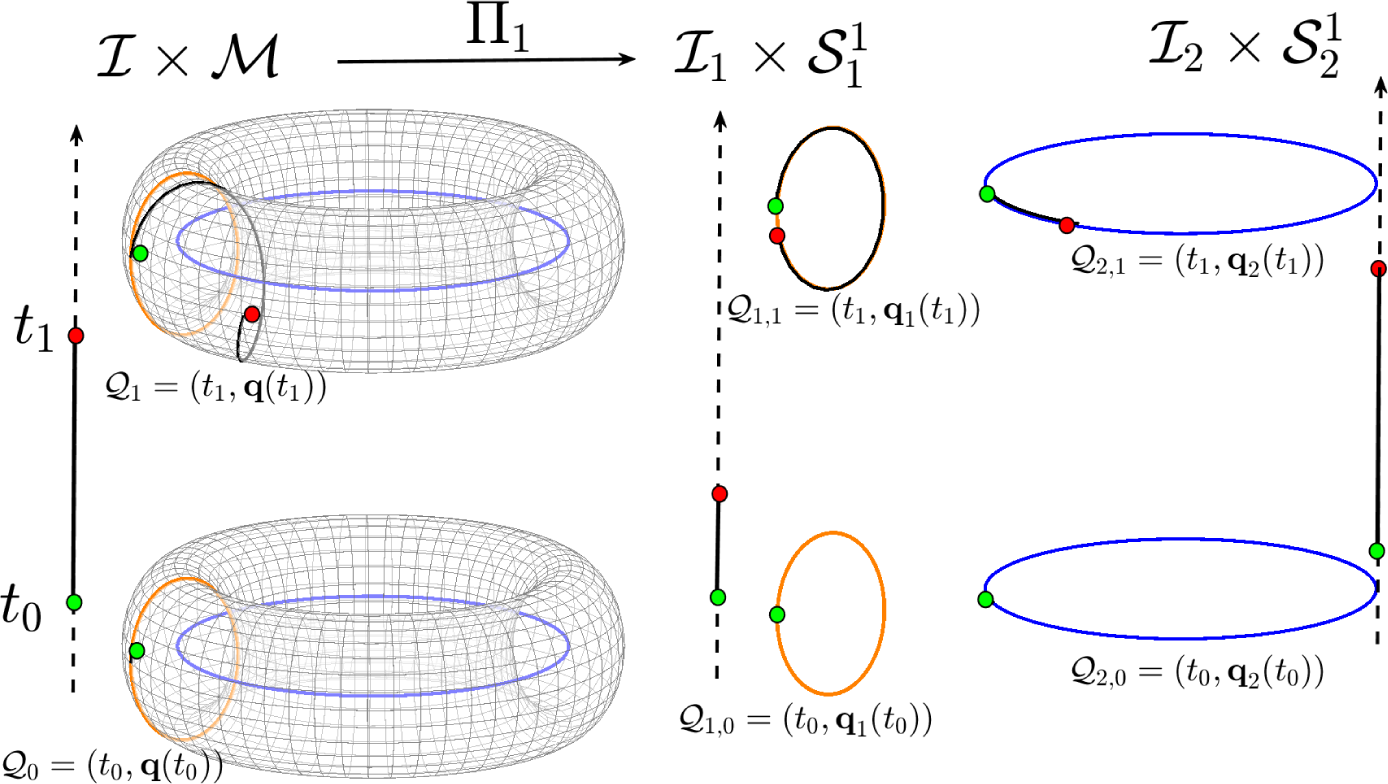
A torus is shown as an example of a possible biological space-time manifold that may be decomposed into submanifolds. The time 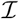 is represented by the vertical arrows. The torus is decomposed into the two circles (sub-manifolds) 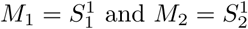. The points 𝒬_0_ at time *t*_0_ are mapped onto the points 𝒬_1,0_ and 𝒬_2,0_ while the point 𝒬_1_ at time *t*_1_ is mapped onto the points 𝒬_1,1_ and 𝒬_2,1_ in the two space-time submanifolds 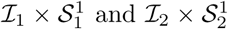. The trajectory on the torus is then mapped onto two different trajectories on the two circles from the initial point (green circle) to the final point (red circle).

#### 2.1.1 Example: decomposing the torus

An example of manifold decomposition is given by the torus, embedded in ℝ^3^ defined as

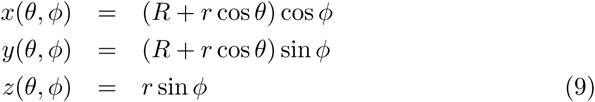

which is decomposed into the two sub-manifolds 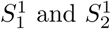, given by

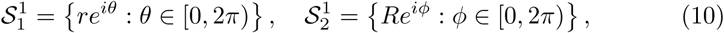

where, *θ* identifies the poloidal direction, *φ* identifies the toroidal direction, *R* the major radius (distance from the center of the torus), and *r < R* is the minor radius of the torus. Therefore, the two biological space-time are given by 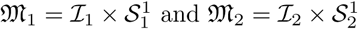. A trajectory Γ on the torus, identified by the equations

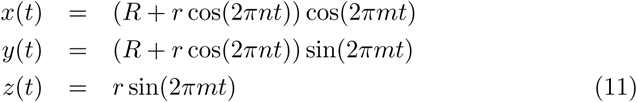

where *t* ∈ [0,1], is then decomposed onto two trajectories

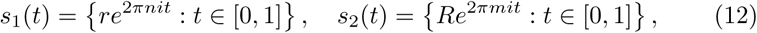

where *n* and *m* are the winding numbers associated with the poloidal and toroidal directions, respectively. In this particular example, the magnitude of the velocity and the acceleration of the projected motion on the two sub-manifolds are given by

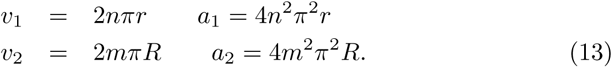

which implies *a_2_ > a_1_* for the same value of *n* and *m*. The position, the velocity and the acceleration on the torus are projected onto the two subspaces in which the dynamics is identified by different values for the position, velocity and acceleration.

The example of the torus illustrates how trajectories on submanifolds (e.g. different biological processes) with different accelerations may be combined and interpreted as a single trajectory on a larger manifold, and vice versa. It must be noted that in this decomposition, the time *t* in the submanifolds is a parameter along the curve and not the biological time. In fact, as we will see in the next section, because the accelerations in the two submanifolds are different we expect time to flow at different rates in each submanifold.

## 3 Dilation and contraction of time

The fact that the Maxwell’s equations are invariant under the Lorentz transformation implies that any inertial observer will measure the light moving at the same constant speed *c*. Moreover, the Lorentz transformations define an invariant quantity, called *proper time*, which represents a space-time interval which assumes the same value for any inertial (non-accelerating) observer. In physics, the presence of fundamental equations such Maxwell’s equations, helps us in understanding the type of coordinate transformations that leave the equations unchanged and can be used to define invariant quantities. Here, we deal with a biological space-time for which the existence of fundamental equations, coordinate transformations, and corresponding invariants, are unknown.

Although many investigators have posed the question of whether or not governing laws exist in biology [Wagner, 2017, Brandon, 1997, Ruse, 1970], we contend this question remains unanswered. Here we assume and postulate the existence of such “biological invariants”, which we note may not have a physical analog. Moreover, it will be reductive to ignore or to assume that fundamental laws of physics will not hold in biological processes. Therefore, in this section we assume the validity of the theory of Special Relativity in biological processes and we investigate its implications by studying the dynamics of a massless point along a trajectory on the manifold related to a biological process. In particular, we consider its dynamics observed by two frames of reference whose coordinates are related by the Lorentz transformations and we investigate the dilation and contraction of biological time resulting from accelerated motions along the trajectory. This approach is used to model and to explore different rates of aging.

We consider two frames of reference *S* and *S′* in the flat Minkowski spacetime *M*^4^ = ℝ × ℝ^3^. In general, the motion of a particle is along a curvilinear trajectory, and the two frames of reference can be in a relative motion like the one represented in Fig. (3a). However, for simplicity, we assume a relative motion with a constant relative velocity *v* along the *x*-direction. We denote with (*t, x, y, z*) and (*t′, x’, y’, z’*) an event recorded by *S* and *S′*, respectively. These two events are linked by the Lorentz transformations

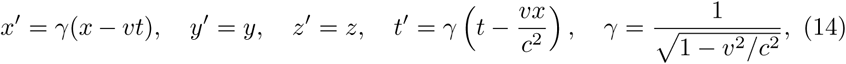

where *γ* > 1 is the Lorentz factor and *c* is the speed of light. It is straightforward to show that these transformations leave invariant the space-time interval *dτ^2^* = *c^2^dt^2^* – *dx^2^* – *dy^2^* – *dz^2^* also called *proper time*. The invariance of the proper time leads to the well known phenomenon of *time dilation*

**Figure 3:**
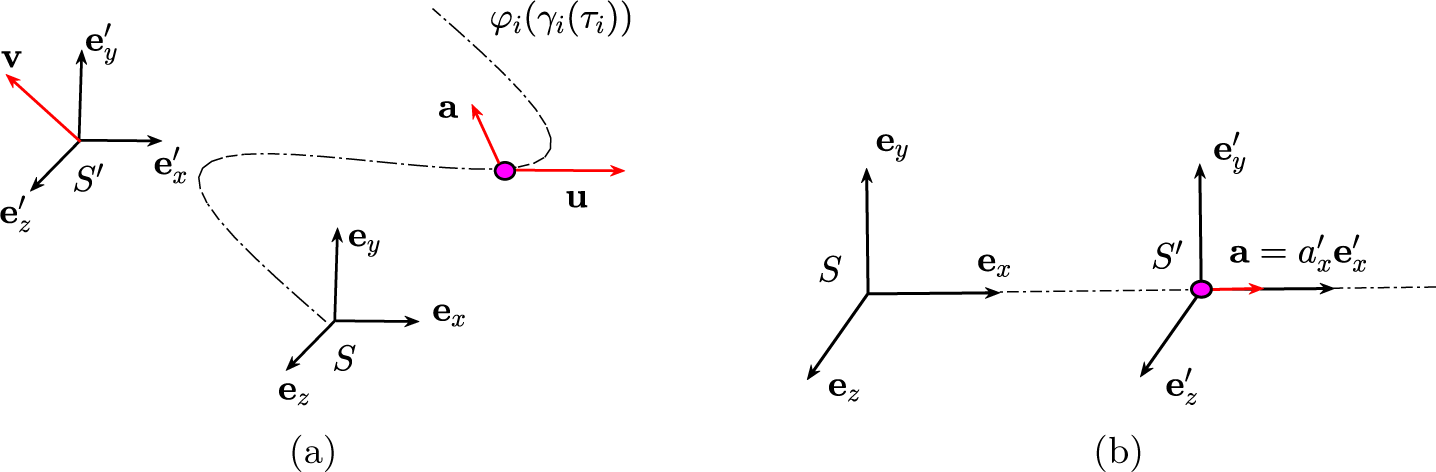
a) The particle (purple dot) is moving along a trajectory with velocity ***u*** and acceleration ***a***. b) The frame of reference *S′* is moving at constant velocity ***v*** away from *S*: the components of the velocity and the acceleration of the particle in these two frames of references are related by Eqs. (16) and (17).

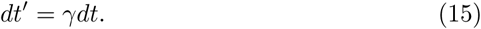

The time interval *dt′* measured from an observer in motion is dilated by the factor *γ* and we say that a moving clock runs slower. It has to be remarked that this effect is reciprocal when both frames of reference are inertial and hence, there is a relative motion at constant speed between them. In that case, no preferred frame of reference exists and each observer can argue that the other one is moving. For this reason, we do consider accelerating frames of reference in which this ambiguity is solved.

By differentiating Eqs. (14) we obtain the transformations for the velocity

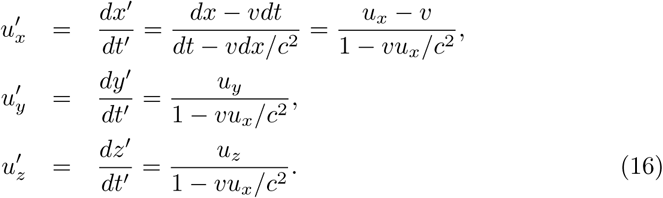

where *u_x_ = dx/dt, u_x_* = *dx/dt* and *u_x_* = *dx/dt* are the velocities along the corresponding direction. An additional differentiation leads to the transformations of the acceleration

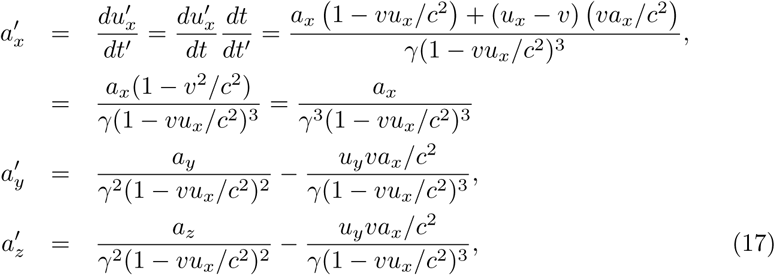

where we have evaluated *dt′/dt* by considering the last of Eqs. (14) and we used the definition of the Lorentz factor *γ*. The sets of Eqs. (16) and Eqs. (17) provide the connection between the velocity and the acceleration of an object measured by an observer in *S* and another observer in *S′*.

We now consider the particular case in which a point is moving along a straight line oriented along the *x*-axis with a constant acceleration 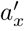 (Fig. (3b)). We assume *S′* to be located at the position of the particle and therefore, its velocity is the same as the velocity of the particle and it increases in time. Hence, by setting *v* = *u_x_* the *x*-component of the acceleration 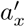 becomes

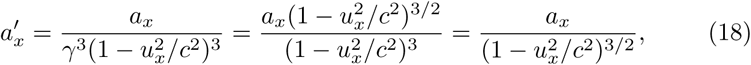

and the acceleration in *S* is given by

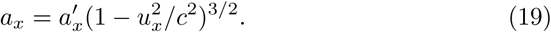

By integrating the above equation with respect to time and by imposing *v*(0) = 0 we obtain the velocity

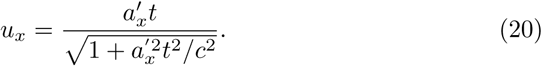

By integrating once again with respect to time and by setting 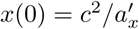 we obtain the position

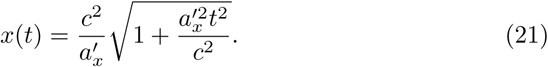

The time *t′* of the accelerated particle, and hence in the accelerated frame of reference, is given by integrating Eq. (15)

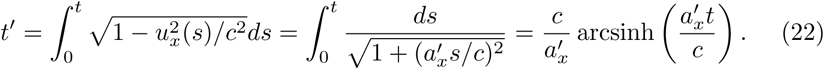

This equation tells us how much time *t′* elapses in the accelerated frame of reference *S′*, in terms of the time *t* elapsed in the inertial frame of reference *S*. This function represents the time component *τ_i_(t)* of the projection map Π*i*, introduced in Eq. (8).

In the case of an accelerating frame of reference, the phenomenon of time dilation is not reciprocal and the observer in the frame of reference *S′* who experiences the constant acceleration cannot argue that the other observer is moving and he is still. Therefore, the time *t′* of the accelerating frame of reference measured from the observer at rest in the frame of reference *S* appears to run slower according to Eq. (22). In what follows we will apply this idea to few biological processes always assuming that the time dilation is described by Eq. (22).

### 3.1 Biological age

Many markers have been used to define biological age. In our modeling framework, biological time is defined as the rate at which biological processes take place, as measured against chronological time. Therefore, our model characterizes one’s biological age by the degree of dilation or contraction of time resulting from the acceleration or deceleration of biological processes that are associated with biological age.

One marker of biological age is the methlyation state of the genome, composed of varying degrees of hyper- or hypomethlyated states [Bocklandt et al., 2011, Horvath, 2013]. The “epigenome” is a component of gene regulation and has been proposed to contain information relevant to the overall state of the biological system [Jenkinson et al., 2017]. Linear age-related epigenetic drift has been associated with cancer incidence [Curtius et al., 2016], and statistical methods have been proposed to identify deviation from time linearity in epigenetic aging, although without a theoretical rationale to describe—or predict—the nature of the nonlinearity [Snir et al., 2016].

In order to explain the process of changing methylation states with age, and the use of methylation state as a surrogate marker of biological age with our model, we first consider the trajectory on a manifold which describes the methylation process of a living person and we assume that this trajectory is mapped onto a straight line in the Minkowski space. We now consider one frame of reference *S* at rest in the origin of the trajectory and one frame of reference *S^’^* that is moving at constant acceleration 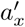 for which the time *t′* is given by Eq. (22). We then assume that the frame of reference slows down to an acceleration 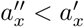 for which the time is *t″* and it is given by Eq. (22). We define the rate of methylation *dM/dt′* and *dM/dt″* when the frame of reference moves with acceleration equal to 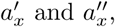 respectively and we have

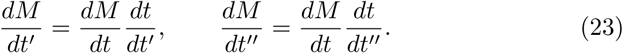

If 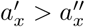 it can be shown that *dt′/dt < dt″/dt*, and therefore, we conclude that

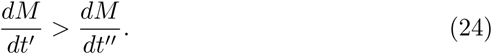

The slowing down of the rate of methylation is therefore interpreted as a decelerating frame of reference. We note that this model does not assume any specific functional form for the methylation trajectory. In fact, during the lifetime of a person, Eqn. (23) permits the acceleration or deceleration of age-related changes to the methylation state, which can be accentuated or modified in the context of cancer.

## 4 Examples of space-time dynamics in biology

In this section we connect our model to different rates of aging between individuals, and discuss examples of precipitous or protracted aging within an individual.

### 4.1 Aging

Aging is defined as the functional and structural decline of an organism, resulting in an increasing risk of disease, impairment and mortality over time. At molecular and cellular level several hallmarks of aging have been proposed to define common characteristics of aging in mammals (Lopez-Otin, Blasco, Kroemer, Cell 2013). These hallmarks include: genomic instability, telomere attrition, epigenetic alterations, loss of proteostasis, altered nutrient-sensing, mitochondrial dysfunction, cellular senescence, stem cell exhaustion and altered cell-cell communication. These processes are interdependent and influenced by cell microenvironmental cues.

Ultimately, the rate of age-related decline varies depending on how genetic variation, environmental exposure and lifestyle factors impact these mechanisms. Consequently, age, when measured chronologically, is often not a reliable indicator of the body’s rate of decline or physiological breakdown. Over the years, the idea of quantifying the “biological age” based on biomarkers for cellular and systemic changes that accompany the aging process have been explored [Levine, 2013], although without a rigorous mathematical treatment of biological age. Major efforts have been made to dissect the individual contribution of each of these hallmarks and factors on aging, but the major challenge remains how to determine how their interconnectedness as a whole impacts the aging dynamics. Here we connect concepts in our mathematical model to biological and chronological aging. We note that for the sake of clarity, we will refer to aging as precipitous (faster than chronological aging) or protracted (slower) in order to clearly associate the terms “accelerated” and “decelerated” to quantities in our modeling framework.

As we derived in the previous section, an unambiguous time dilation effect requires an accelerating frame of reference. We now imagine a range of range of accelerations representing a distribution within a population between the values *a_min_* and *a_max_* (green region in Fig. (4a)). The dashed and dotted curves corresponds to Eq. (22) by using the values *a_min_* and *a_max_*, respectively. Any acceleration within this range will correspond to a curve in the green region. An interval of time Δ*t* on the *x*-axes corresponds to a reduced interval of time Δ*t′* on the *y*-axis and therefore, the ticking of the second clock is slowed down. The existence of a distribution, or range of values of acceleration is necessary to capture both features of time dilation, (green region in Fig. (4a)) and time contraction, when the value of the acceleration is less than *a_min_* (pink region). Moreover, values of the acceleration much larger than *a_max_* can be used to define the arrest of biological time such as in a hibernating state or cryogenic freezing (blue region). The case of zero acceleration is given by Eq. (15) and is not considered in this model context.

**Figure 4:**
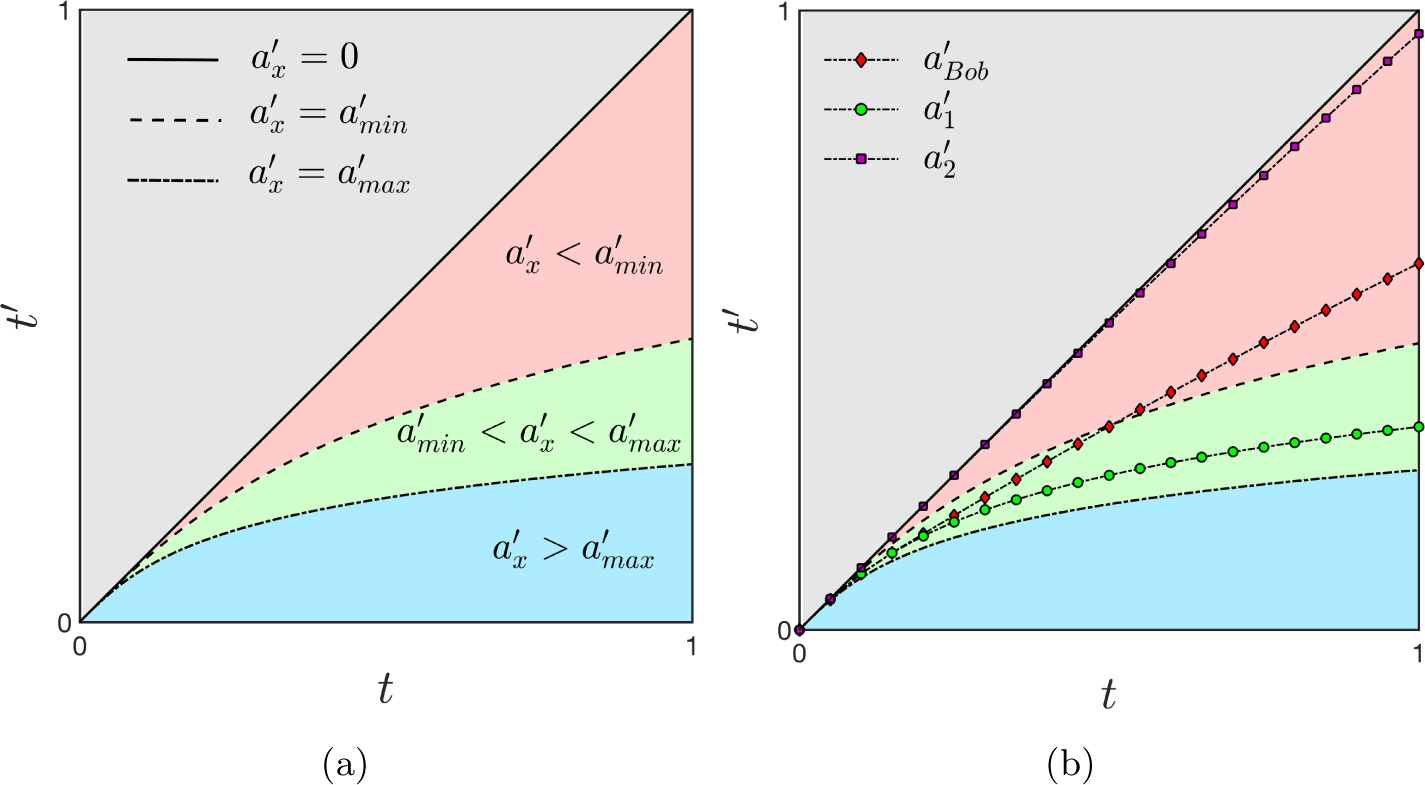
a) For an accelerating frame of reference a time interval At Δ*t*= 1 corresponds to shorter Δt′ with a consequent dilation of time. The green region represents a distribution of accelerations associated with a average range of accelerations that is used as a common reference for accelerated and decelerated processes. Smaller values of the acceleration are related to time contraction (pink region, 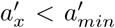) while large values are associated with time dilation (blue region, 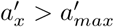).b) In the mental illustration of the lifetime of Bob, a deceleration will bend Bob’s trajectory (black line with red diamonds markers) out of the green region with a corresponding contraction of time.

#### 4.1.1 Alice and Bob

To illustrate the contraction and dilation of time in biological space-time (i.e. on a submanifold) and how they can be related to the presence of a disease, we consider the following thought experiment involving Alice and Bob, two individuals with different rates of aging. For simplicity, we assume that their dynamics occur on a flat torus *τ^2^* defined as the Cartesian ℝ^2^ plane under the identifications (*x, y*) ∼ (*x + 2π, y)* ∼ (*x, y + 2π*). The opposite edges *x, x + 2π* and *y, y + 2π* of the domain are identified and therefore, they must be interpreted as the same point (see red and blue edges in Fig. (5)). In other words, the trajectory that comes out from the right edge of the plane, will come in from the left edge and the same applies from the upper and lower part of the plane. In this case, the manifold is trivially mapped into ℝ^2^ with the identity map, and hence the dynamics on the manifold corresponds to the dynamics in the Minkowski space-time.

**Figure 5:**
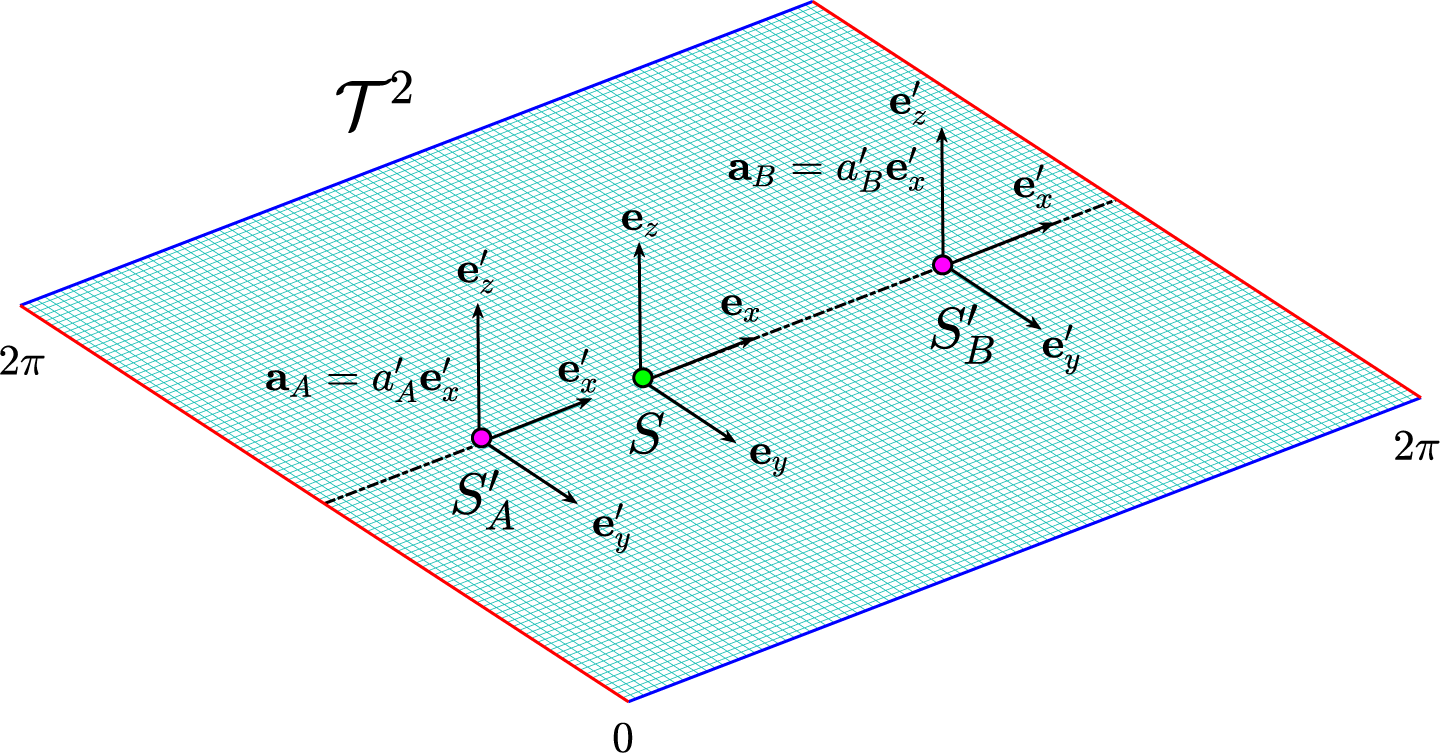
The flat torus *τ^2^* in which the dynamics of Alice 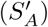 and Bob 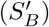 occur along the straight dotted line. The green and the purple dots represent their birth and their actual position on the manifold. The effect of time dilation is more evident for Alice whose frame of reference moves at higher acceleration.

The point on the manifold corresponding to the birth of Alice and Bob is indicated in Figure (5) by the green circle in which we assume to place a frame of reference *S* identified by the three unit vectors (*e_x_, e_y_, e_z_*). The dynamics of Alice is represented by the motion of a purple point in which we imagine to place a frame of reference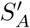. We also assume that the point is moving along a straight line with constant acceleration given by 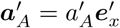. The same conditions apply to Bob, but in this case the frame of reference is 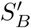 and the particle is moving with a constant acceleration 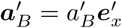 (see Fig. (5)).

We now can apply the results obtained in the previous section to infer that the time in 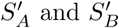 are given by Eq. (22)

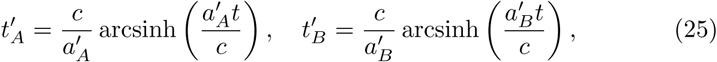

Assuming the acceleration of Alice to lie within the range [*a_min_, a_max_*], her proper time 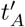 lies in the green region (see Fig. (4a). On the other hand, Bob’s frame of reference is moving with acceleration 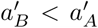 and its proper time 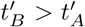 deviates from the green region towards an older biological age in the pink region. With respect to Alice, Bob is moving with a slower acceleration and hence, his clock is running faster, resulting in a time contraction with respect to Alice’s clock. If instead we assume 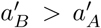, Bob’s proper time 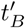 will be larger than 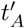 and it deviates from the green towards the blue region. In this case Bob’s clock will run slow with a consequent more evident effect of time dilation. Health, therefore, could be generally compared between individuals by determining their accelerations as compared to a frame of reference at rest.

We now consider only Bob and we assume his frame of reference 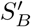 to have an acceleration 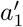 within the range 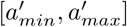. Suddenly, his acceleration is decreasing to the value 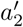 and we therefore want to know what will happen to its proper time *t′*. In particular we hypothesize the acceleration to change as follows

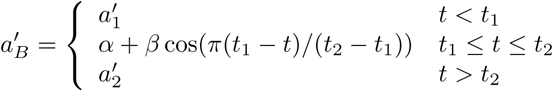

where 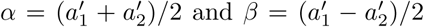. By solving Eqs. (20) and (22) we obtain the time *t′* for the decelerating frame of reference shown in Fig. (4b): the line marked with green circles and the line with violet squared markers represent the trajectory corresponding to the accelerations 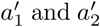, respectively. However, due to the deceleration, the line marked with red diamonds will not follow the line with the green circles but it will move outside the green region.

The deceleration of Bob’s frame of reference is reflected to a contraction of time while a larger time dilation is a consequence of an increasing accelerating frame of reference.

#### 4.2 Cancer as inducer of precipitous aging

Major pathologies, such as cancer, diabetes, cardiovascular disorders and neurodegenerative diseases have an impact on aging. Cancer and chemotherapy in particular are known to accelerate the aging process [3]. As described above, aging involves multiple complex changes at molecular and cellular level that lead to decline in physiologic reserve capacity across virtually all organ systems [13]. In cancer, accumulation of cellular damage aggravates the hallmarks of aging and accelerates aging. In cancer patients, the trajectory of decline worsen not only because of the direct physiologic insult inflicted by cancer, but also because of the injuries induced by anti-cancer therapies, such chemotherapy or radiation, to systems that maintain physiologic reserves [8, 7]. In this context, there is a dose-dependent effect, whereby the more intensive the treatment of cancer or more vulnerable and frail the physiologic state, the steeper the decline in physiologic reserves [13, 12]. Below, we use these two variables, intensity of therapy and frailty, to illustrate our model of accelerated aging.

It is not surprising that the two cancer populations most affected by precipitous aging caused by cancer are survivors of childhood cancer who are typically exposed to intensive multi-agent therapy at a young age [Henderson et al., 2014, Ness et al., 2015] and adult patients who undergo hematopoietic cell transplantation (HCT) for refractory hematologic malignancies [M et al., 2016]. These two scenarios can be explained by our model as follows. The early onset of advanced biological age in childhood cancers corresponds to a rapid contraction of time that deviates away from the green region, similar to Bob’s modified trajectory, but with an earlier onset (see Fig. (4b)). The case of allogeneic HCT, in which case the transplanted cells come from another person, corresponds to a discontinuity in biological space-time. The recipient’s trajectory on the hematopoietic manifold and/or the biological age is reset to that of the donor. If the biological ages of the donor and recipient of the HCT do not agree, the recipient will experience an abrupt and persistent change in the rate of biological aging corresponding to the discontinuity in both biological space and time.

Frailty, characterized by a cluster of measurements of physical states, is the best described measure of aging in a population, and identifies individuals who are highly vulnerable to adverse health outcomes and premature mortality. Although frailty is not a perfect corollary with biological age, it is a measure of abnormal aging at a population level. In long-term survivors of childhood cancer (median age 33 years [range 18–50 years]), the prevalence of frailty [Rockwood and Mitnitski, 2011] has been shown to be comparable to that reported among adults greater than 65 years of age [Ness et al., 2013]. Comparable high rates have been reported in adult survivors of HCT [M et al., 2016]. In fact, frail HCT recipients have a three-fold increased risk of subsequent mortality compared with the non-frail counterparts,[M et al., 2016] which is similar to the downstream consequences of frailty seen in the general population such as adverse health outcomes [Rockwood and Mitnitski, 2011], and early mortality [Fried et al., 2001, Hogan et al., 2003]). From the lens of our model, frailty can be associated with a biological age and therefore defined as a threshold in time 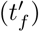 so that the onset of frailty in the cancer population 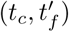 will appear sooner relative to the general population 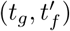 where *t_c_ < t_g_*.

##### 4.2.1 Chronic inflammation

The normal process of aging is associated chronic low grade inflammation and with cumulative oxidative stress, independently of disease. Inflammation and oxidative stress are critical responses in host defense and injury repair and are essential for normal body functions. However, with advanced age there is a loss of sensitivity in the injury-repair cycle leading to persistent chronic inflammation, and a natural decline in the endogenous anti-oxidant capacity leading to cumulative oxidative stress [Mittal et al., 2014]. These two process are interdependent, and contribute to the hallmarks of aging, influencing telomere length, mitochondrial function, epigenetics and stem cell self-renewal.

Chronic inflammation and oxidative stress are also common underlying factors of age-associated diseases. Inflammation, particularly chronic low grade inflammation, has been found to contribute to the initiation and progression of multiple age-related pathologies such as type II Diabetes, Alzheimer’s disease, cardiovascular disease and cancer [Mantovani et al., 2008]. In addition, chronic diseases, and in particular cancer, elicit and promote an inflammatory tumor microenvironment that increases cancer fitness. Therefore, chronic inflammation has been suggested to underlie and accelerate biological aging [Fougère et al., 2017], in particular when associated with disease. The concept of “inflamm-aging”, inflammation-associated aging, can be used to provide a systemic perspective of biological aging in the human population [Franceschi et al., 2007].

On the level of tissue homeostasis, the best characterized example is hematopoietic aging associated with chronic inflammatory signaling in the bone marrow microenvironment. In fact, the “age” of a young hematopoietic stem cell can be “reprogrammed” when transplanted into an aged or inflammatory environment [Kovtonyuk et al., 2016], highlighting the impact of “inflamm-aging” and the plasticity of molecular clocks. Aging is associated with clonal hematopoiesis and accumulation of mutations in hematopoietic progenitors [Steensma et al., 2015], likely due to the underlying inflammation in the aged bone marrow niche. Indeed, an inflamed bone marrow can induce pre-leukemic conditions in mice similar to those occurring in elderly patients [Wang et al., 2014, Dong et al., 2016]. Our model can be applied to measure how microenvironmental cues, such inflammation, impact aging of hematopoietic cells. Changes in the biological age of the cell can be represented as a periodicity in the biological manifold resulting in an abrupt change in the dilation or contraction of time.

#### 4.3 Protracted aging

In contrast to precipitous biological aging which corresponds to the contraction of biological time, protracted aging corresponds to the dilation of biological time. This can be illustrated by prolonged periods of near zero biological activity, for instance in freezing conditions or hibernation which is part of a continuum of biological and metabolic states [van Breukelen and Martin, 2015]. How can this situation be explained in a relativistic framework? We imagine a *cryogenic*-manifold mapped onto a Minkowski space. If in this space the frame of reference is moving with a high acceleration in the the direction of motion then the interval of time in its frame of reference tends to zero: a frozen cell corresponds to a frame of reference moving at very *large acceleration relative to its unfrozen state* on this particular biological manifold. Figure (4) shows the limiting case of a near perfectly frozen (cryogenic) biological process (blue region), which would correspond to an extreme dilation of time.

## 5 Discussion

Here we have investigated a mathematical model of biological aging. We define a biological space-time mathematically as the Cartesian product of manifolds and sub-manifolds and apply the principles of special relativity to compare different rates of biological aging. We illustrate the concepts of precipitous and protracted aging as the contraction and dilation of time with a mental illustration comparing the lifespans of two individuals, Bob and Alice. This analogy provides a framework to compare the rates of aging between individuals by determining their rate of acceleration as compared to a common frame of reference.

Other groups have proposed a framework for biological time to explain biological rhythms and other oscillating or biological processes that repeat periodically[Bailly et al., 2011]. Our biological space-time model is more general, and is aimed towards explaining nonlinear biological phenomena that are not necessarily periodic or oscillating, although these can be considered special cases of more generalized trajectories. In complement to Noble’s hypothesis that there exist no privileged level of causation in biology[Noble, 2012], we suggest that the multiple scales of biology may be interpreted as trajectories on distinct manifolds that may be combined and coupled to each other. Although generalizations of relativity and space-time have been proposed, here we adapt these mathematical structures in order to interpret trajectories on manifolds to represent biological processes [O’Neill, 1983].

Our model introduces many questions and suggests novel hypotheses. In particular we considered the implication of the Lorentz transformations and, by thought experiments, we give a possible explanation to time contraction and dilation in terms of decelerating and accelerating frame of reference. However, when considering biological processes there can be different coordinates transformations which may define a *biological invariant*, which could be a quantity, or point in biological space-time. A biological invariant in a biological space-time may allow us to characterize and define the mathematical laws that govern biological space-time and whether or not the laws remain constant under transformations within that space. The characterization of such an invariant quantity could have profound consequences for our understanding of biological space-time and the potential role of relativity in this space to model the process of aging. A limitation of our approach is the degree of uncertainty in the appropriateness of the use of the Lorentz transformation in a biological context, which requires further study to confirm or falsify.

A novel concept which naturally follows from our approach is the notion of *force* in a biological context. A biological force may be a generalization of a physical force in a biological context which changes the dynamics of a particle during its dynamics on a manifold. We may hypothesize then, that accelerated dynamics on the manifold may be the effect of the resultant of biological forces acting on the particle. For example, cancer and chemotherapy may be interpret as forces which combine and have a resultant which may change the dynamics of the point on a manifold with a consequent impact on the rate of aging.

The shape of the manifolds which characterize biological space-time also affect our notion of distance, which requires the knowledge of the metric tensor of the manifold. Although a manifold is defined to be locally Euclidean, two biological processes or objects that are sufficiently different from each other(in space-time) may not be measured by using the conventional definition of Euclidean distance; ex. distances on Earth are locally linear but not globally. The degree to which we must redefine our notion of a *metric* in the biological space-time may also affect our ability to interpret long-time dynamical behaviors of biological systems. We therefore hypothesize that the underlying geometry of the manifold, or equivalently, the elements and structure of the metric tensor, may provide valuable insights into the biological process.

## 6 Summary

In summary, we have proposed a mathematical framework that may be used to define a biological space-time. We use this as a tool to model and study different rates of biological aging based on the concept of the dilation or contraction of time due to the acceleration or deceleration of biological processes relative to a common frame of reference. We discuss some examples of biological processes that illustrate these concepts, and discuss implications and novel hypotheses that are generated by this model.

## 7 Acknowledgements

Research reported in this publication was supported by the National Cancer Institute of the National Institutes of Health under award number P30CA033572. The content is solely the responsibility of the authors and does not necessarily represent the official views of the National Institutes of Health. We thank Leo D. Wang for a critical reading of the manuscript, and Jacob G. Scott for discussions that inspired and motivated this work. DM would like to thank G. Bocca for useful discussions.

